# Rapid genomic evolution in *Brassica rapa* with bumblebee selection in experimental evolution

**DOI:** 10.1101/2021.12.02.470896

**Authors:** Léa Frachon, Florian P. Schiestl

## Abstract

**Background:** Insect pollinators shape rapid phenotypic evolution of traits related to floral attractiveness and plant reproductive success. However, the underlying genomic changes remain largely unknown despite their importance in predicting adaptive responses to natural or to artificial selection. Based on a nine-generation experimental evolution study with fast cycling *Brassica rapa* plants adapting to bumblebees, we investigate the genomic evolution associated with the previously observed parallel phenotypic evolution. In this current evolve and resequencing (E&R) study, we conduct a genomic scan of the allele frequency changes along the genome in bumblebee-pollinated and hand-pollinated plants and perform a genomic principal component analysis (PCA).

**Results:** We highlight rapid genomic evolution associated with the observed phenotypic evolution mediated by bumblebees. Controlling for genetic drift, we observe significant changes in allelic frequencies at multiple loci. However, this pattern differs according to the replicate of bumblebee-pollinated plants, suggesting putative non-parallel genomic evolution. Finally, our study underlines an increase in genomic differentiation implying the putative involvement of multiple loci in short-term pollinator adaptation.

**Conclusions:** Overall, our study enhances our understanding of the complex interactions between pollinator and plants, providing a steppingstone towards unravelling the genetic basis of plant genomic adaptation to biotic factors in the environment.

## Background

Pollinator insects are important selective agents for wild- and crop plant species due to their essential role in the reproduction of most flowering plants [1]. While a decline of pollinator insects has been detected in different geographical regions and insect families [2, 3, 4], the understanding of the adaptive potential of plants to such changes remains in its infancy. Plant adaptation to pollinators typically involves traits associated with flower attractiveness such as (1) flower morphology [5, 6, 7], flower colour [8, 9], flower scent [10, 11, 12], and (2) traits associated with mating system like herkogamy [13, 14] or selfing [15, 16]. While most studies assessed the result of long-term evolutionary adaptation to pollinators, tracking the adaptive processes across generations remains scarce. Both the resurrection approach in natural populations, growing seeds from different generations together, or experimental evolution studies, applying the same selective pressure for multiple generations, can bridge this gap. For instance, using a resurrection approach, Thomann et al. (17) observed phenological and reproductive trait changes over 18 years in *Adonis annua* plants in response to the loss of wild bees. While this approach benefits from ecological realism in natural populations, it makes it difficult to differentiate the effect of the factor of interest from other factors such as climate, also shaping plant evolution. Gervasi and Schiestl [12] performed experimental evolution with fast-cycling *Brassica rapa* plants evolving with different pollinators and under controlled conditions, to identify the evolutionary response to pollinator-mediated selection. They showed, within nine generations of experimental evolution, rapid plant adaptation to bumblebee pollination in phenotypic traits, such as floral volatiles, UV reflection and plant height. However, while these evolved traits have also been shown to carry substantial heritability in this study system [18, 19], the genomic changes underlying these rapid plant phenotypic changes are still unknown.

In the current context of pollinator decline and the associated changes in pollinator communities, understanding the genetic architecture involved in plant response to pollinators is essential to understand their adaptative potential in changing environments [20, 21, 22]. Molecular genetic studies have uncovered the molecular and genetic bases of several traits involved in pollination and pollinator attractiveness such as selfing [23], pollination syndromes [24, 8, 25, 26, 27], nectar [28,29] and volatiles [30, 31, 32, 33, 34]. However, insects use a combination of signals (shape, colour, scent) and rewards for identifying suitable flowers leading to plant adaptation based on multiple traits [35]. For instance, honest signals (signals associated with reward) and pollination syndromes (convergent evolution of specific signal combinations selected by pollinators) are good examples of evolution of multiple traits. In a context of rapid environmental changes, genetic correlation among traits may allow the synchronous response of different phenotypic traits to varying patterns of selection [36, 37, 19]. While essential to predict the adaptive potential of plants to pollinators and enable breeding of crop that are more attractive to pollinators, we are still in the beginning of understanding the genetic basis involved in the adaptative response to pollinators. By combining experiment evolution and next generation sequencing (NGS), evolve and resequencing (E&R) studies have proven their ability to unravel the genomic basis involved in rapid evolution [38, 39, 40].

Here, based on previous experimental evolution performed by Gervasi and Schiestl [12] with outcrossing fast-cycling *Brassica rapa* plants, we tracked the genomic changes involved in the adaptative response of plants to bumblebee selection compared to hand pollinated control plants in an E&R study (**Figure 1**). Gervasi and Schiestl [12] initiated their experimental evolution with 108 full sib seed families in the greenhouse. Over nine generations of selection, they grew plants in three isolated replicates (36 plants per replicate). The selection was mediated by three pollination treatments during the experiment: bumblebee (*Bombus terrestris*) pollination, hoverfly- (*Episyrphus balteatus*), and hand pollination. Overall, they observed parallel phenotypic evolution with bumblebee-pollinated plants evolving taller, more fragrant flowers, and being more attractive to bumblebees. In our E&R study, we re-sequenced plants from two replicates in bumblebee and hand-pollination treatments. We performed a genome scan of allele frequency changes and a genomic principal component analysis to observe putative genomic evolution shaped by bumblebees.

**Figure 1.**
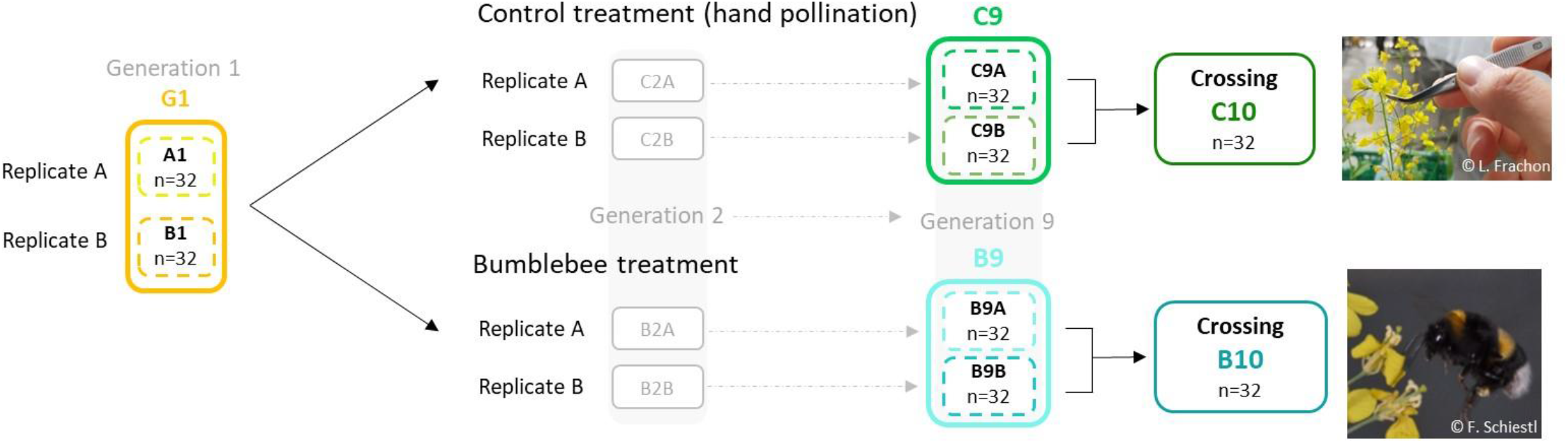
Experimental evolution design. Partial experimental evolution design from Gervasi and Schiestl [12] from the first generation (G1) to the ninth generation in control (C9) and bumblebee treatment (B9). The tenth generation was obtained by inter-replicate crossing (C10 and B10). Population of *Brassica rapa* fast cycling used for seedling sequencing are coloured. The abbreviation and the colours assigned to the populations are unchanged along the manuscript. The number of seedlings (n) sequenced per population is indicated. Images: courtesy of F. Schiestl and L. Frachon.

## Results

### Genomic changes during experimental evolution

In our study, we observed allele frequency changes (Δ*h*) over nine generations in both bumblebee and control treatments and between two replicates within treatments (**Figure 2**). Based on 10’000 genetic drift simulations, we defined significant changes (colored dots in **Figure 2**) as observed changes in allele frequency outside the drift simulation ranges (upper and lower grey lines in **Figure 2**). The allele frequency changes varied substantially between treatments and between replicates within treatments. For instance, we found 32 SNPs with strong (Δ*h* > 0.5) and significant (fdr < 0.05) changes in one of the replicates of the bumblebee treatment (replicate A). In contrast, we observed only 4 SNPs with significant allele frequency changes in the other replicate in bumblebee treatment (replicate B). In the control (hand-pollinated) treatment, there were 3 SNPs with significant allele frequency changes in one replicate (B) and zero SNPs with allele frequency changes in the other replicate (A). In replicate A, we found a 42-fold increase in the number of SNPs displaying significant allele frequency changes (fdr > 0.05) with the bumblebee treatment compared to the control treatment, regardless of its strength i.e. considering all the value of Δ*h* (4 SNPs with significant changes in the control treatment, and 171 SNPs in the bumblebee treatment, **Figure 2**). However, no difference in the number of SNPs involved in significant allele frequency changes between bumblebee and control treatment were observed in replicate B (21 SNPs with significant changes in the control, and 19 SNPs in the bumblebee treatment considering fdr < 0.05, considering all value of Δ*h*, **Figure 2**).

**Figure 2.**
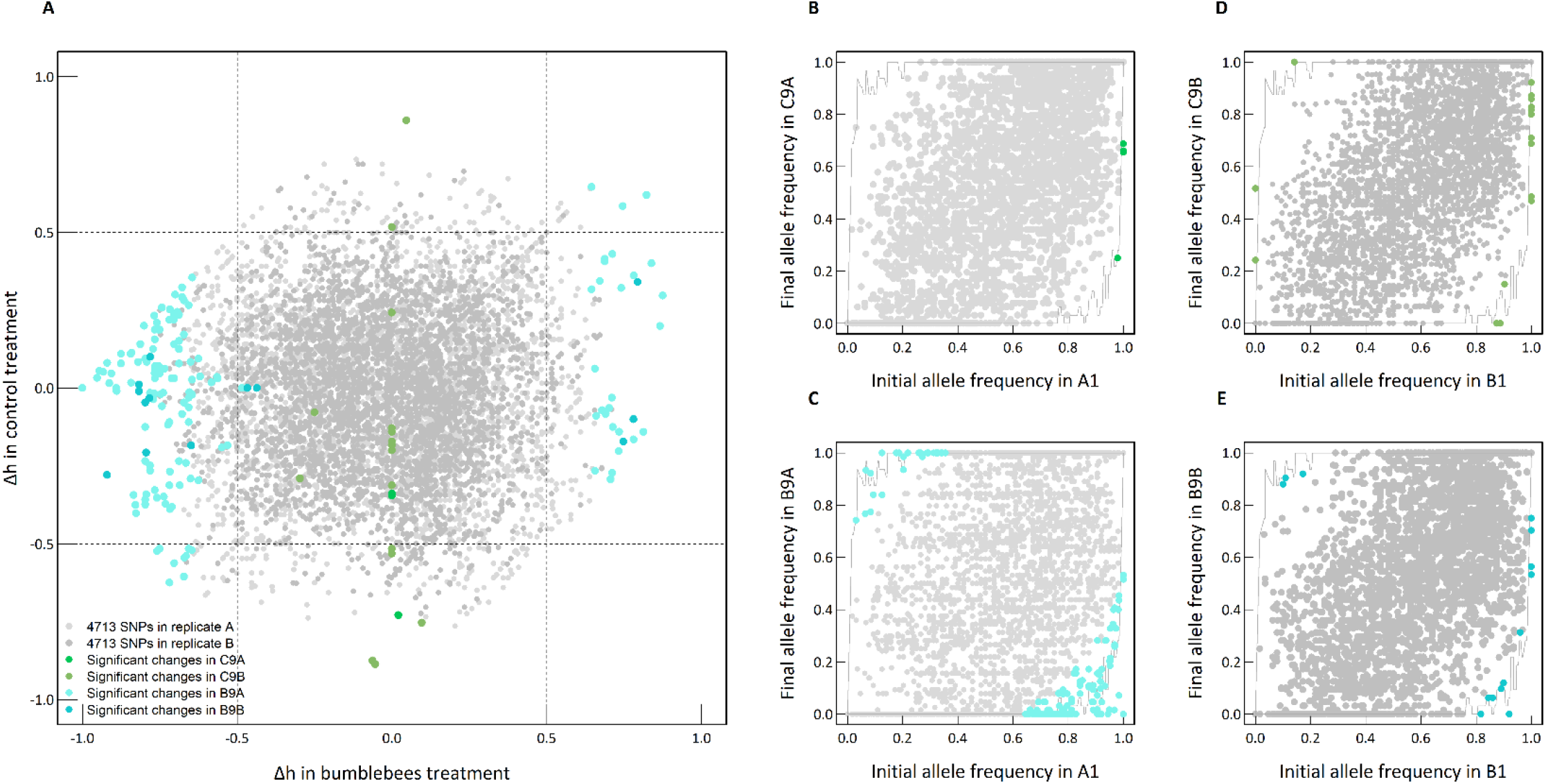
Allele frequency changes during experimental evolution. (**A**) Comparison of the allele frequency changes (Δh) between the bumblebee treatment (*x-axis*) and the control treatment (*y-axis*). The grey dots represent the 4’713 SNPs. Replicate A is in light grey, replicate B in darker grey. The significant changes are highlighted in blue and green (see the legend in the figure for details). Comparison of initial (first generation) and final (ninth generation) allele frequencies in the control treatment for both replicate A **(B)** and replicate B **(D)**, and in the bumblebee treatment for both replicate A **(C)** and replicate B **(E)**. The grey dots represent the non-significant changes in allele frequencies between generations. The grey solid lines indicate the maximum (upper line) or minimum (lower line) of final simulated allele frequencies obtained by 10’000 simulations of random genetic drift (over nine generations, N_e_=16). The color dots represent important changes with *fdr* < 0.05.

As expected, the evolutionary process is associated with an increase of linkage disequilibrium in the bumblebee treatment, but also in the control treatment. In both treatments, we observed a slower decay of the median linkage disequilibrium in the ninth generation compared to the first generation (**Figure S3A**). Within the bumblebee treatment, we observed a slower decay of LD in replicate A than in B as expected with the observed stronger allele frequency changes (**Figure S3A**). The decays of LD in inter-replicate crossing were slower in both treatments compared to the ninth generation (**Figure S3A**). Moreover, we observed a decrease of the number of haplotype blocks, *i*.*e*., genomic regions with strongly linked alleles, over nine generations (from 786 and 780 haplotype blocks for replicate A and B respectively in the first generation, to 563 and 648 haplotype blocks for replicate A and B respectively in the ninth generation of the control treatment, and 478 and 600 for replicate A and B respectively in the ninth generation of the bumblebee treatment) associated with an increase of their length (**Figure S3BC, Table S1**). We observed 743 and 696 haplotype blocks in control and bumblebee treatments respectively in the inter-replicate crossing (**Figure S3BC, Table S1**). It is worth noting that the moderate density of genetic markers (18 SNPs per Mb in mean among the 10 chromosomes, **Figure S1**) suggests that the estimate of LD and haplotype length may be slightly underestimated, although this has no effect on the trends observed.

Overall, the SNPs involved in the evolutive responses are independent between replicates, and between treatments (**Figure 3**). For instance, among the 171 SNPs with significant allele frequency changes in replicate A in bumblebee treatment, 164 SNPs are unique of this population, whereas 5 SNPs are shared with the control treatment in replicate B, and 3 SNPs are shared with the bumblebee treatment in replicate B (**Figure 3**).

**Figure 3.**
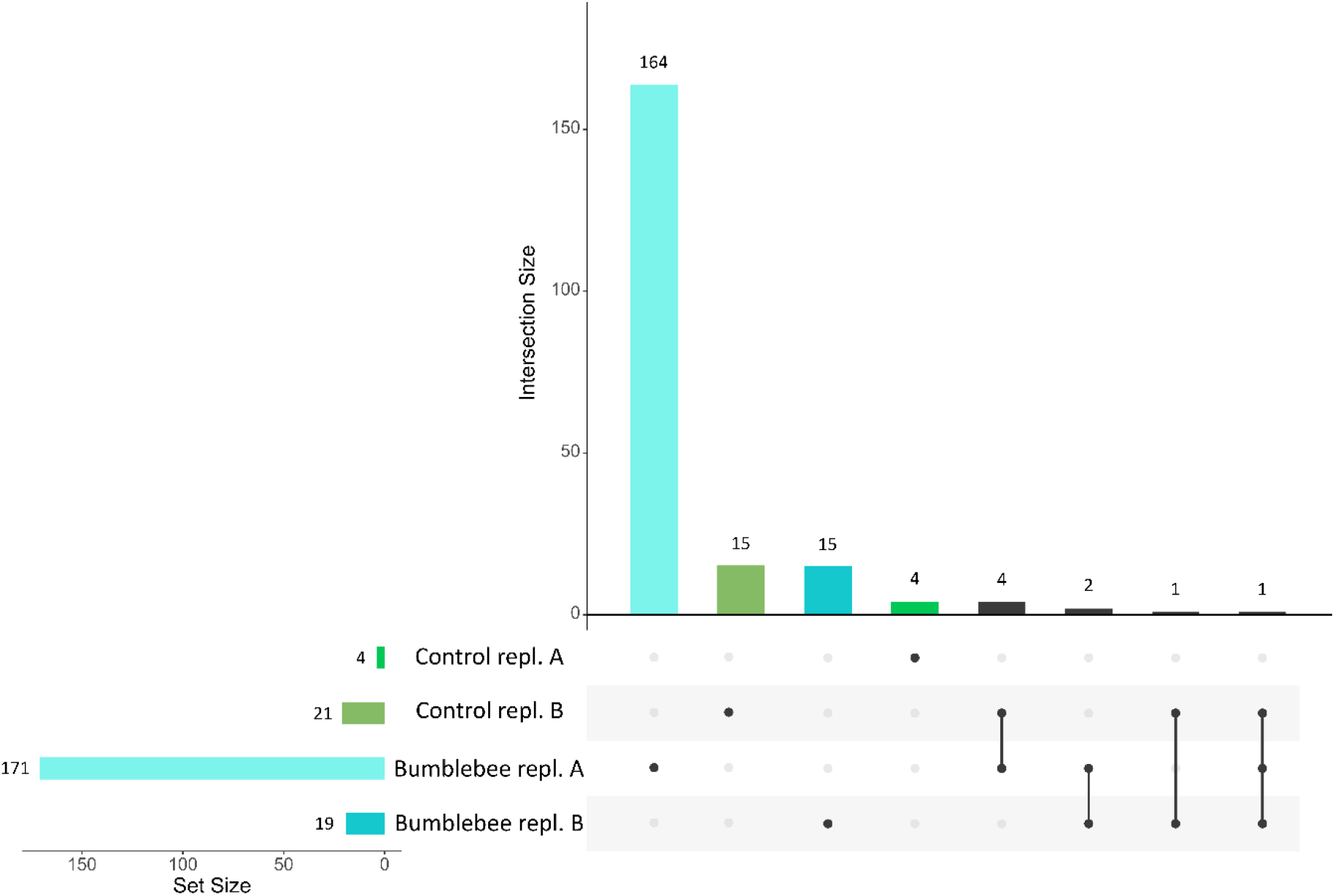
Intersection of genomic variants under selection between treatment and replication. The UpSet plot illustrates the genomic variants involved in the evolution of *Brassica rapa* in control and bumblebee treatment for both replicates A and B. On the left (Set Size), the number of significant genomic variants involved in evolutionary response of *B. rapa* in different populations is indicated. The dots indicate that the genomic variants are involved in the evolutionary response only in a particular population, while the vertical lines indicate that the genomic variants are involved in the evolutionary processes in several populations. The number of genomic variants involved in unique population or in different populations is indicated in the upper part of the figure.

### Identity of candidate genes underlying genomic evolution in bumblebee treatment

After retrieving the annotated genes around 4kb (2kb upstream, 2kb downstream) for 187 SNPs with significant allele frequency changes in the bumblebee treatment (fdr < 0.05), we obtained a list of 171 candidate genes (**Dataset1**). Briefly, we found genes with biological functions that may be involved in the emission of volatiles (ABC transporter G), in the regulation of flowering time (B3), or in shoot architecture (UCH). In addition, some genes are involved in the response to different biotic and abiotic stresses (METACASPASE-2, PUB, Pum, PAT8, BON). Finally, some candidate genes do not have a known biological function. However, due to a putative underestimate of LD and haplotype length (see previous section), we may not have been able to correctly identify all the candidate genes involved in this evolutionary process. **Genome-wide population structure**. To assess the relatedness among individuals across generation and treatments, we performed a genomic principal component analysis (PCA) on the full set of SNPs (4’713 SNPs). We observed a structuration of our samples determined by populations *i*.*e*. generations, treatments, and replicates (**Figure 4**). Along the six first principal components (PCs) axis, explaining 56.07% of the total genomic variance (**Figure 4A**), all individuals from the two replicates of generation one were grouped together, and individuals from the ninth generations were clearly separated from the first generation (**Figure 4**). In this genomic space, individuals from the inter-replicate crossing fell between those from the scatterplot formed by replicates A and B of generation nine for both treatments (**Figure 4**). Interestingly, in the genomic space created by the first 6 principal components explaining the most genetic variance, the individuals resulting from selection by bumblebees were always more scattered than those from the control treatment or the first generation.

**Figure 4.**
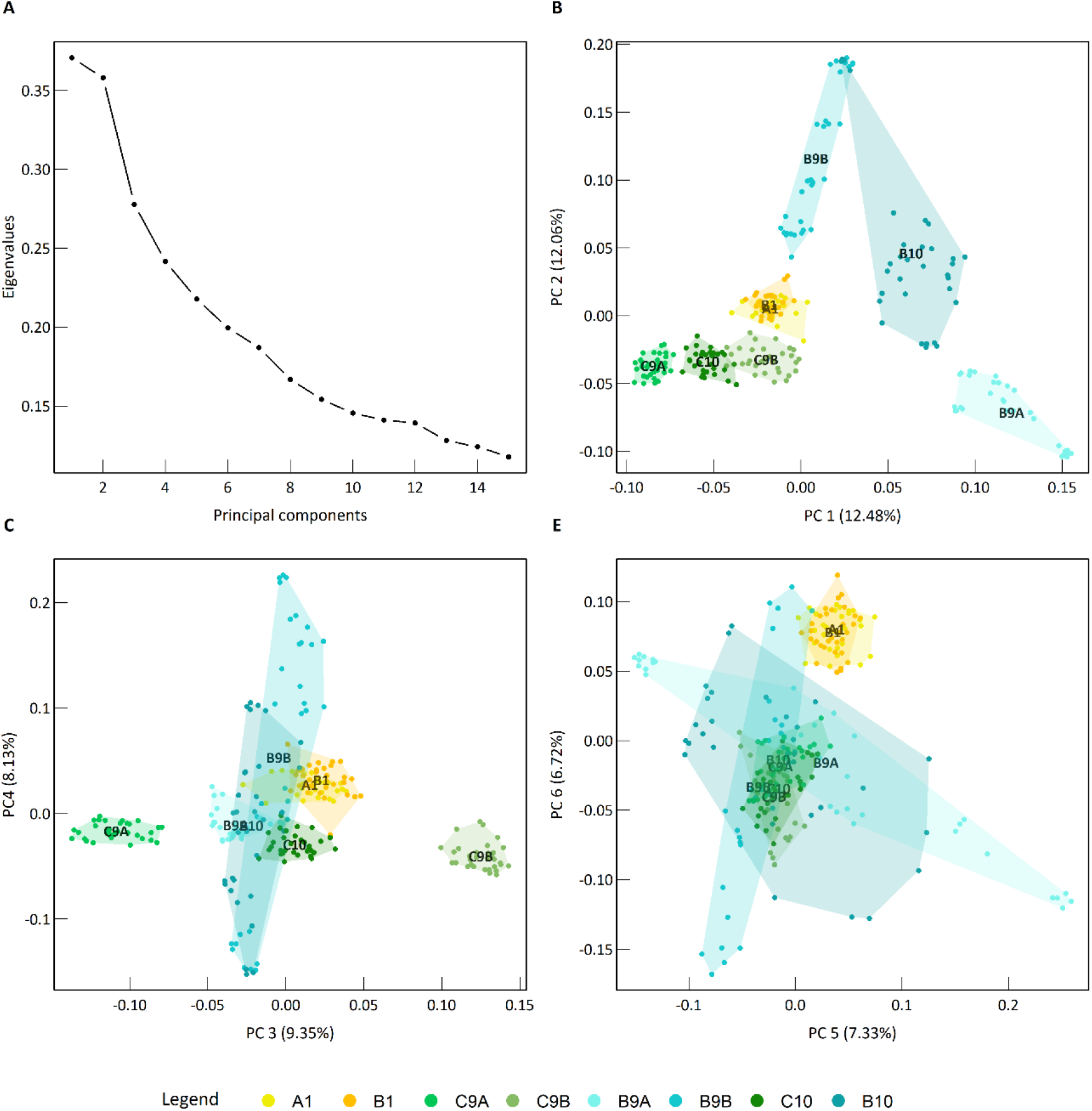
Genome-wide population structure. (A) Eigenvalues of the 15 principal components of the genomic PCA based on 4713 SNPs. **(B-D)** Position of the 256 individuals in the genomic space from the principal component analysis (PCA) performed on their genotypes (GT). The label of the population is shown on their centroid. Polygons linking outer points of the scatter plot are displayed for all populations. The legend colours are indicated in the bottom of the figure.

## Discussion

Understanding pollinator-mediated evolutionary genomics of flowering plants remains an important challenge in biodiversity conservation and crop improvement. Here, we screened for the genomic consequences of biotic selection in an E&R study by sequencing genome-wide SNP markers in *Brassica rapa* plant individuals before and after nine generations of selection by bumblebees, and under random hand-pollination. The previous experimental evolution has shown, at the phenotypic level, that this primarily outcrossing plant rapidly adapts to specific pollinators [12]. For instance, Gervasi and Schiestl [12] observed taller plants with an increase in total scent emission per flower in bumblebee-pollinated plants. The pattern of phenotypic evolution was similar between replicates from the same pollination treatment. In this E&R study, we documented the putative signature of directional selection driven by bumblebee pollinators with significant allele frequency changes at several loci. Moreover, we have shown different pattern of genomic evolution between replicates, and an increase of genomic differentiation among individuals.

In agreement with the demonstrated phenotypic evolution in the fast-cycling *Brassica rapa* experimental system [12], we have shown genomic evolution across nine generations. The here documented changes in allele frequencies, the increase of linkage disequilibrium and the number of haplotype blocks, underscore the importance of pollinators in shaping plant rapid genomic evolution. The limited number of individuals (36 per replicate and per treatment) and replicates (2 replicates) could impact the evolutionary processes and the identification of the underlying genomic bases. Indeed, small population size can lead to an increase in the effect of genetic drift, and a low number of replicates to a poor estimate of loci under selection [41]. However, at the phenotypic level, the observed differences among the three replicates were smaller than those observed among treatments, indicating that pollinator selection played a larger role than random drift [12]. Moreover, in our study, we accounted for genetic drift bias and identified changes in allele frequency that were greater than those expected from 10’000 genetic drift simulations. While we cannot completely exclude some effects of genetic drift, this alone cannot explain the patterns observed. Moreover, some significant candidate genes identified in our E&R study have biological functions in line with the phenotypic evolution of traits observed by Gervasi and Schiestl [12]. For instance, while the previous experimental evolution study highlighted an increase in scent emission and plant height driven by bumblebee pollination, we identified candidate genes who, or at least their family, are known to be involved in emission of volatile organic compounds, like ABC transporter G family (ABCG35, ABCG38, ABCG1, ABCB19, [42, 43]), in shoot architecture like Ubiquitin C-terminal hydrolase (UCH [44]), or in growth and development like AUXIN-RESPONSIVE PROTEIN-RELATED, or TREHALOSE-PHOSPHATE PHOSPHATASE E-RELATED [45]. The U-box protein is also an interesting candidate gene due to its involvement in different developmental and physiological processes such as self-incompatibility, defence, or abiotic stress response [46]. Moreover, we found significant allele frequency changes associated with flowering time like a B3 transcription regulator [47] or UCH [48]. While Gervasi & Schiestl [12] did not observe any difference in flowering time between bumblebee and control treatment in the last generation, flowering time is an important phenological trait involved in plant-pollinator matching [49, 50]. It is therefore possible that pollinators induce flower phenological shift, which was not detected in the phenotypic data, but picked up in the genomic changes as changes influencing flowering time. Overall, our study highlighted the potential involvement of multiple loci in rapid adaptation to bumblebees, which agrees with studies highlighting such a genetic architecture underlying floral evolution [51, 26,52, 37,53]. However, further analysis using higher quality of sequencing and the characterization of associated phenotypic and phenological traits (allowing a genome-wide association approach) and functional validation of the genes are required to draw more solid conclusions about the genetic basis of evolutionary changes induced by bumblebee selection.

In addition, we found that the extent of genomic changes observed during the evolutionary processes was different between replicates suggesting non-parallel genomic evolution [54]. While Gervasi and Schiestl [12] have shown convergent (or parallel) phenotypic evolution for the bumblebee treatment, our results indicated that only one replicate exhibited several loci with significant changes in allele frequency (i.e. outside the range of drift simulations). Such a convergent phenotypic evolution associated with non-parallel genomic evolution has been observed in artificial selection of crops [55, 56] and in natural populations. For instance, a recent study highlighted different genomic regions underpinning the evolutionary convergence of herbicide resistance in blackgrass among different populations [57]. Likewise, in natural populations of *Senecio lautus*, similar phenotypic variation was reported to be regulated by variation in genomic space across populations in dune-headland coastal habitats [58]. However, the low number of replicates in our study and the significant changes in allele frequencies observed in only one replicate do not provide a clear distinction between potential non-parallel evolution and a “two-speed” parallel genomic evolution (*i*.*e*. the second replicate will follow the same evolutionary path in the upcoming generations). Longer experimental evolution, with more replicates and individuals are needed to clearly understand the evolutionary pattern mediated by biotic factors such as bumblebees.

Finally, we observed an increase of genomic variance, within the genomic space created by the main differentiation axes, during experimental evolution mediated by bumblebees [12]. This increase in overall genomic variance was observed among individuals in both bumblebee-pollinated replicates (B9A and B9B), as well as in the inter-replicate crossing (B10). This pattern might be explained by the polygenic model with a weaker selection acting on multiple standing variants (soft sweep) and by multiple loci underlying individual phenotypic trait evolutionary changes. An increased number of studies demonstrate the importance of polygenic adaptation [56, 60, 61, 62] related to the infinitesimal model (reviewed in Barton et al. [63]), where local adaptation is driven by small allele frequency changes in multiple loci. Interestingly, highly polygenic architecture involved in phenotypic evolution could contribute to the maintenance of standing genetic variation, as recently demonstrated in long-term artificial selection on chicken weight [64]. However, deepened analyses are needed using newly developed models to validate the involvement of polygenic architecture in rapid phenotypic evolution [65, 66, 67]. Multiple genes underlying phenotypic variation are widely emphasized in plants with the advances of GWAs [59, 62], however the interplay of evolutionary forces on these genes is still poorly understood and deserves further studies.

## Conclusion

We revealed important genomic changes on multiple loci associated with convergent phenotypic evolution resulting from bumblebee selection in only nine generations. Our study is a first step into the understanding of the complex genomic mechanisms involved in rapid evolutionary adaptation to biotic factors, and we advocate further analyses to understand (1) the genetic architecture underlying phenotypic and phenological variation, (2) pleiotropic effects of quantitative-trait locus in rapid adaptation and (3) the mechanisms behind a maintenance of genetic variance. We also underline the importance of better characterizing the gene functions involved in plant-pollinator interactions. Overall, pollinators constitute complex patterns of selection which deserve more attention for predicting the adaptive responses of wild and crop plant species to their decline.

## Methods

### Plant material and experimental design

*Brassica rapa* (Brassicacea) is an outcrossing plant with genetic self-incompatibility, pollinated by diverse insects such as bumblebees, flies or butterflies [68]. Our study used rapid-cycling *Brassica rapa* plants (Wisconsin Fast Plants™, purchased from Carolina Biological Supply Company, Burlington, USA), selected for its short life cycle of approximately two months from seed to seed in our greenhouse conditions. We used seeds produced by the study of Gervasi and Schiestl [12], performing experimental evolution with bumblebees and control hand pollination. Briefly, starting from 108 full sib seed families, the pollination was carried out over nine generations either by bumblebees (*Bombus terrestris*), hoverflies (*Episyrphus balteatus*) or by random hand cross-pollination (control treatment). This experiment was conducted with 3 isolated replicates (one replicate includes 36 plants) for each treatment. A representative subset of seeds from all pollinated flowers was used for the next generation; the contribution of seeds to the next generation being calculated as 36 divided by the sum of seeds per replicate over all individual seeds. In this study, we focused on bumblebee and hand-pollination treatment, using two most distinct replicates (replicate A and B). We used the offspring of the starting populations and the ninth generation for both treatments. On seed per individual plant were used. In total, we used 32 32 plants from the starting generation (generation 1) in replicate A (called A1), and 32 plants from replicate B (called B1), 32 plants from the ninth generation selected by bumblebees (bumblebee treatment) in replicate A (called B9A) and 32 plants in replicate B (called B9B)and ; 32 plants from the ninth generation of control hand pollination plants (control treatment) in replicate A (called C9A) and replicate B (C9B; **Figure 1**). Finally, we performed crossings between replicates A and B within each treatment (generation 10) yielding 32 individuals from the bumblebee treatment (inter-replicate crossing in bumblebee treatment; here called B10) and 32 individuals from the control treatment (inter-replicate crossing in control treatment; here called C10). This manual crossing is commonly used for reducing the effect of potential inbreed depression on trait changes. Pollen donors and receivers were randomly assigned. Each combination of generation*treatment*replicate is called a population (*e*.*g*. ninth generation, treatment bumblebees, replicate A called B9A is a population). A total of 256 seeds from these 8 populations (first, ninth and tenth generation) were sown out in a phytotron (first generation in 2017 and ninth generation as well as the inter-replicate crossing in 2019) and the leaf tissue of each plant was collected for DNA extraction and whole genomic sequencing. The study conforms to the institutional, national, and international regulations.

### DNA extraction and genomic characterization

Because leaf tissue was collected in 2017 for the first generation and 2019 for the last generations, we adapted the collection storage (drying vs freezing). Leaf material from the first generation was dried in vacuum at 40 °C for 20 hours, and leaf material from the ninth and tenth generation was stored in -80°C. A high molecular weight DNA extraction (average DNA concentration of 48 ng/μL, LGC extraction protocol) and library preparation for genotyping-by-sequencing (restriction enzyme MsII, insert size mean range ∼215bp) was performed by the LGC Genomics group Berlin. Samples were sequenced with Illumina NextSeq 500 V2 sequencer using 150 paired-end reads; the alignment of our samples was performed with BWA version 0.7.12 against the reference genome sequence of *Brassica rapa* FPsc v1.3, Phytozome release 12 (https://phytozome.jgi.doe.gov/pz/portal.html) by the LGC Genomics group Berlin. The variant discovery and the genotyping were realized using Freebayes v1.0.2-16 with the following parameters by the LGC Genomic Group Berlin: --min-base-quality 10 –min-supporting-allele-qsum 10 –read-mismatch-limit 3 –min-coverage 5 –no-indels –min-alternate-count 4 –exclude-unobserved-genotypes –genotype-qualities –ploidy 2 or 3 –no-mnps –no-complex –mismatch-base-quality-threshold 10. We then performed a quality trimming on the 10 chromosomes *i*.*e*. by discarded unassigned scaffolds from the chromosomes using vcftools [69], by removing SNPs with missing data in more than 5% of the individuals (function –max-missing 0.95, *i*.*e*. genotype calls had to be present for at least 243 samples out of 256 for a SNP to be included in the downstream analysis), and retaining only bi-allelic SNPs with a minimum average Phred quality score of 15 (function –minGQ 15, **Figure S2DE**). We removed SNPs with a maximum average read depth of 100 (function –max-meanDP 100, **Figure S2BC**), and discarded SNPs with a minor allele frequency (MAF) lower than 0.1 (function –maf 0.1, **Figure S2A**). The final dataset contained 4’713 SNPs in ∼ 283Mb genome size (https://phytozome.jgi.doe.gov/pz/portal.html).

### Allele frequency changes and genetic drift simulation

The allele frequencies of the reference allele for the 4’713 SNPs were estimated within each 8 populations using vcftools (function –freq, [64]). To control potential genetic drift during the nine generations of evolution, we simulated random final allele frequencies 10’000-fold for different ranges of initial allele frequencies (from 0 to 1 by an interval window of 0.01). The simulations were performed using the R environment package “learnPopGen” (function “drift.selection”, [70]) over eight transitions between generations (*i*.*e*. from the first generation to the ninth generation) considering 32 individuals within each population for an effective size (N_e_) of 16 (*i*.*e*. individuals contributing to the next generation, see details of experimental evolution in Gervasi and Schiestl [12]), and considering an equal fitness for each individual. For each SNP along the genome, we assigned their initial allele frequency value (AF_initial_) to the range estimated by the final allele frequency simulations (10’000 values). Finally, the observed final allele frequency (AF_final_) was compared to 10’000 simulated allele frequency values (AF_simulated_) to estimate a p-value for each SNP using the following equations:

1. For a decrease of reference allelic frequency *i*.*e*. (AF_initial_ – AF_final_) > 0, *pvalue* = (number of simulations with AF_simulated_ ≥ AF_final_)/10’000
2. For an increase of reference allelic frequency *i*.*e*. (AF_initial_ – AF_final_) < 0, *pvalue* = (number of simulations with AF_simulated_ ≤ AF_final_)/10’000
3. For (AF_initial_ – AF_final_) = 0, *pvalue* = 1

With AF_simulated_ = simulated final allele frequency, AF_initial_ = initial allele frequency from the reference allele (first generation), and AF_final_ = observed final allele frequency for the reference allele (ninth generation). The pvalues were controlled for the False Discovery Rate (fdr) of 5% using Benjamini-Hochberg method implemented in R environment [71].

Finally, we estimated the allele frequency changes (Δ*h*) from the reference allele according to the equation (1) for both bumblebee and control treatments:

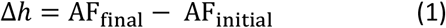

Where Δ*h* is the allelic frequency change between the first and the ninth generation, AF_initial_ is the initial observed allele frequency, and AF_final_ is the observed final allele frequency at the ninth generation.

### Linkage disequilibrium (LD) and haplotype block structure evolution

During the selective process, an increase of the linkage disequilibrium *i*.*e*. the non-random association of alleles between loci, is expected in genomic regions strongly under selection. First, we calculated pairwise linkage disequilibrium (LD) among all set of SNPs measured by the correlation coefficient between SNP pair r^2^ within each chromosomes using VCFtools (function –geno-r2) for each 8 populations containing 32 individuals. The associated median LD was then estimated and plotted. Finally, to understand whether changes in median LD are due to random allelic associations along the genome, or aggregated in genomic regions under selection, we calculated the haplotype blocks in each population using plink1.9 with the following parameters: --blocks no-pheno-req –maf 0.07 –blocks-max-kb 200. This method estimates the length of these blocks with “strong LD” considering the allelic association D’ metrics between 0.7 and 0.98 according to Gabriel et al. [72].

### Candidate genes

We identified 171 candidate genes associated with 187 SNPs with significant allele frequency changes (fdr < 0.05) during bumblebee selection. Because the median linkage disequilibrium in bumblebee treatment is ∼4kb, we retrieved the annotated genes around 4kb (2 kb upstream and 2 kb downstream) for 187 SNPs and extracted the gene description using phytozome.jgi.doe.gov.

### Genome-wide variance changes

The genomic variance among individuals within each population was estimated using principal component analysis (PCA) on scaled and centered genotype data (pcadapt package in R environment, function pcadapt, [73]). In order to unravel the changes in genomic variance over nine generations, we performed the PCA (1) on the total number of SNPs (4713 SNPs), and (2) on 187 SNPs showing significant allele frequency changes under bumblebee selection in on or another replicate (fdr < 0.05). To compare genomic variance among populations, we estimated the area of the polygon formed by scatter plots of axis 1 and 2 of each population for the full dataset (4713 SNPs) and the subset of 187 SNPs using R package tidyverse and splancs [74, 75]. The relative area is the ratio between the polygon area within each population and the polygon area circumventing all the points of the PCA.

## Declarations

### Ethics approval and consent to participate

The study conforms to the institutional, national, and international regulations.

### Consent for publication

Not applicable.

### Availability of data and material

The datasets used and/or analysed during the current study are available from the corresponding author on reasonable request. The DNA sequences of all samples will be available at National Library for Biotechnology Information (NCBI) database (BioProject PRJNA931117) after acceptance of the manuscript. Reviewers can access to the data using the following link: https://dataview.ncbi.nlm.nih.gov/object/PRJNA931117?reviewer=hocdpu4je8n22ueldu7potnjqn

### Authors’ contributions

L.F. and F.S. designed the research. L.F. performed the DNA extraction and statistical analyses and wrote the manuscript. L.F. and F.S. reviewed and edited the manuscript.

## Acknowledgements

We thank Jörg Vogt for the first-generation DNA extraction, and the LGC Genomic group Berlin for the DNA extraction of other samples, the sequencing and bioinformatic treatment. We thank Cyril Zipfel for discussion regarding candidate genes. We thank Loren Rieseberg and Marius Rösti for their useful comments on a previous version of the draft. Finally, we thank the University of Zürich for their financial support.

