## Supplementary Information for "Rapid genomic evolution in *Brassica rapa* with bumblebee selection in experimental evolution"

**Figure S1. Density of genetic markers for each 10 chromosomes.**

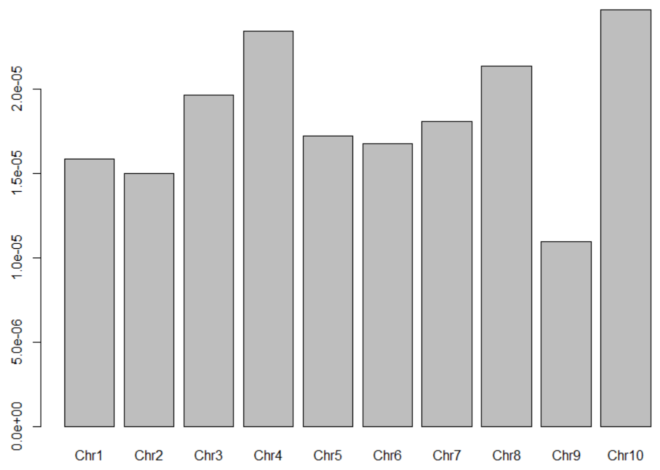

**Figure S2. Distribution of filtered genomic data for our final dataset of 4'713 SNPs. (a) Distribution of the minor allele frequency, (b) distribution of average read depth (DP) per SNPs, (c) distribution of the average read depth (DP) per individual, (d) distribution of the average genotype quality (GQ), (e) different thresholds of the minimum average GQ as a function of the number of SNPs in the final dataset. The red line indicates the chosen value in our study (--minGQ = 15).**

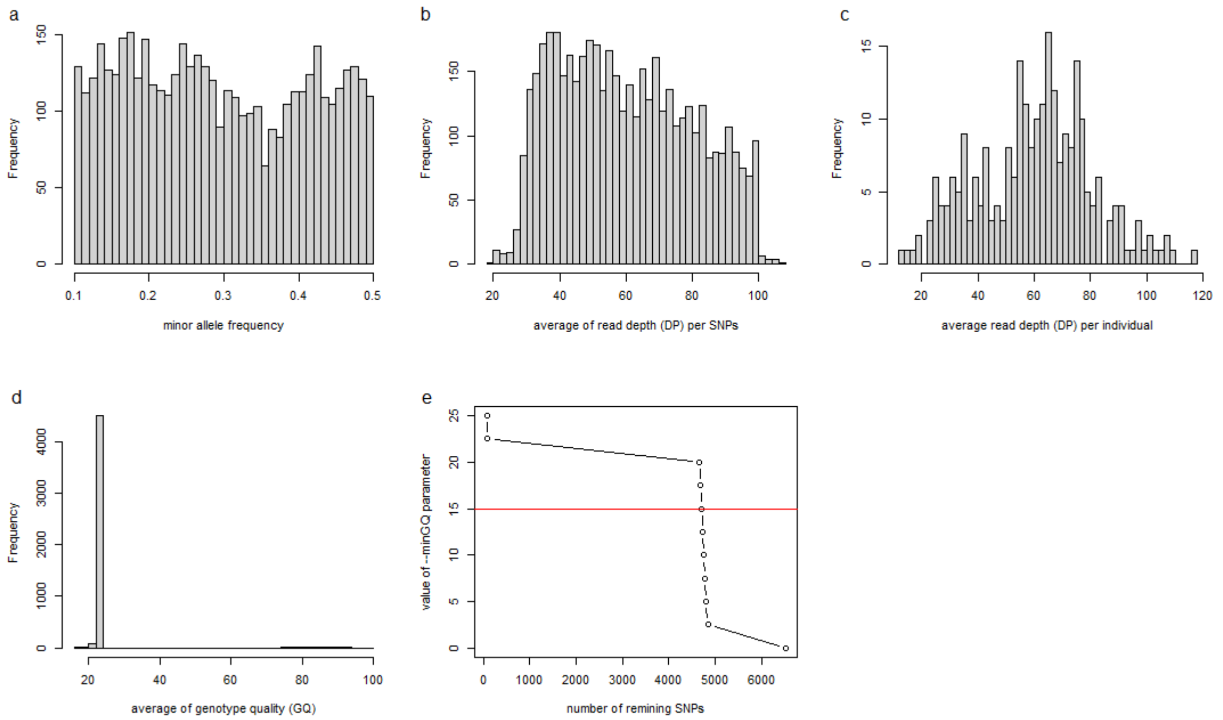

**Figure S3. Linkage disequilibrium and haplotype block structure. (A)** Distribution of the median pairwise linkage disequilibrium ($r^2$) for each population by distance between two SNPs (kb). **(B)** Number of haplotype block calculated within each population **(C)** Average length (kb) of haplotype blocks per population (more details Table S1).

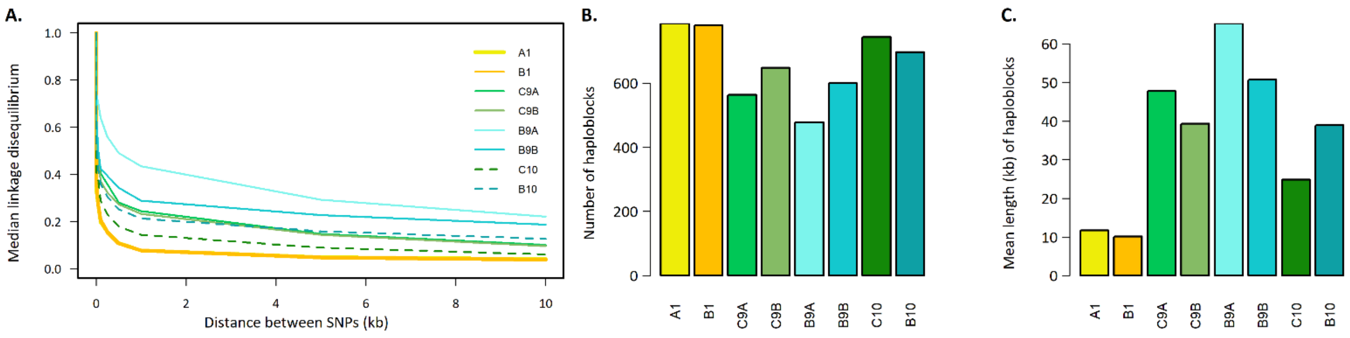


**Table S1. Haplotype blocks.** For each population, the number of haplotype blocks, the mean (+- sd) of the number of SNPs per haplotype block, and the length (in kb) of haplotype blocks.

| | Nber blocks | mean length/block | median length/block | mean nber of SNP/block | median nber of SNP/block |
|---|---|---|---|---|---|
| A1 | 786 | 11.75 | 0.11 | 3.13 | 3 |
| B1 | 780 | 10.14 | 0.1 | 3.06 | 3 |
| C9A | 563 | 47.81 | 0.69 | 4.07 | 3 |
| C9B | 648 | 39.37 | 0.2 | 3.91 | 3 |
| B9A | 478 | 65.33 | 20 | 4.33 | 4 |
| B9B | 600 | 50.71 | 1.13 | 4.12 | 3 |
| C10 | 743 | 24.87 | 0.15 | 3.53 | 3 |
| B10 | 696 | 38.94 | 0.17 | 3.87 | 3 |

**Dataset1. List of candidate genes with significant allele frequency changes during bumblebee selection.** The transcripts are retrieved from phytozome.jgi.doe.gov.
